# *APOE4* drives transcriptional heterogeneity and maladaptive immunometabolic responses of astrocytes

**DOI:** 10.1101/2023.02.06.527204

**Authors:** Sangderk Lee, Holden C. Williams, Amy A. Gorman, Nicholas A. Devanney, Douglas A. Harrison, Adeline E. Walsh, Danielle S. Goulding, Tony Tuck, James L. Schwartz, Diana J. Zajac, Shannon L. Macauley, Steven Estus, TCW Julia, Lance A. Johnson, Josh M. Morganti

## Abstract

Apolipoprotein E4 (APOE4) is the strongest risk allele associated with the development of late onset Alzheimer’s disease (AD). Across the CNS, astrocytes are the predominant expressor of *APOE* while also being critical mediators of neuroinflammation and cerebral metabolism. APOE4 has been consistently linked with dysfunctional inflammation and metabolic processes, yet insights into the molecular constituents driving these responses remain unclear. Utilizing complementary approaches across humanized APOE mice and isogenic human iPSC astrocytes, we demonstrate that ApoE4 alters the astrocyte immunometabolic response to pro-inflammatory stimuli. Our findings show that ApoE4-expressing astrocytes acquire distinct transcriptional repertoires at single-cell and spatially-resolved domains, which are driven in-part by preferential utilization of the cRel transcription factor. Further, inhibiting cRel translocation in ApoE4 astrocytes abrogates inflammatory-induced glycolytic shifts and in tandem mitigates production of multiple pro-inflammatory cytokines. Altogether, our findings elucidate novel cellular underpinnings by which ApoE4 drives maladaptive immunometabolic responses of astrocytes.

## Introduction

Beyond aging alone, carriage of the ε4 allele of the *APOE* gene (APOE4) represents the greatest risk factor for late onset Alzheimer’s disease (AD); individuals expressing a single allele have a threefold increase in risk, while homozygous carriers are up to fifteen times more likely to develop AD. In the central nervous system (CNS), ApoE is expressed predominantly by astrocytes, to which ApoE4 has been demonstrated to alter multiple kinetics associated with amyloid-beta (Aβ)/tau production, clearance, and aggregation. Furthermore, there is growing appreciation for the diverse roles of astrocytic ApoE beyond AD-related proteinopathies ^1^.

Astrocytes are CNS-resident glia that maintain tight regulation of a variety of functions associated with development, homeostasis, and metabolic support ^2–4^. However, in disease settings, astrocytes’ close proximity to all other CNS resident cell types positions them as central mediators and potent propagators of neuroinflammation ^5^. This connectivity has been demonstrated to drive complex reactive states of astrocytes in a variety of neuroinflammatory settings ^6–13^. Problematically, ApoE4 has been connected with disrupting a variety of homeostatic astrocyte functions as well as exaggerating degenerative reactivity associated with neuroinflammation and cerebral metabolism ^14^. Recent work has demonstrated that selective removal of the ε4 allele from astrocytes is sufficient to abrogate several disease-associated neuroinflammatory- and reactive astrocyte correlates ^15,16^. Further, recent insights have highlighted unique astrocyte heterogeneity in response to systemically-induced neuroinflammation ^17^. Despite consistent findings correlating ApoE4-driven astrocyte dysfunction knowledge surrounding the underlying mechanisms and whether there are allele-dependent effects of ApoE variants upon transcriptional and metabolic heterogeneity remains limited.

In the current study, we focused on examining astrocyte-specific responses to pro-inflammatory challenge to ascertain the molecular underpinnings associated with ApoE4-driven immunometabolic responses. We describe a novel interaction of ApoE4-expressing astrocytes, relative to ApoE3, to preferentially acquire transcriptionally distinct heterogeneity and metabolic inflexibility, which are attributable to utilization of the canonical NF-κB transcription factor cRel. We demonstrate that disruption of cRel activity blunts inflammatory-induced and ApoE4-exacerbated alterations at the level of single cell and spatial transcriptome, metabolome, and proteome, thereby highlighting a translatable therapeutic pathway.

## Results

### ApoE4 drives transcriptional heterogeneity of astrocytes following systemic LPS administration

While *APOE* has been shown to modulate peripheral and myeloid immune response, the effects of E4 carriage on astrocyte-specific outcomes to an innate immune challenge are not well-defined. Thus, we first examined the effect of ApoE4 upon astrocyte transcriptional responses using single cell RNA sequencing (scRNAseq) 24hrs following systemic lipopolysaccharides-induced (LPS; 5mg/kg) neuroinflammation. Dimensional reduction and visualization via UMAP revealed 12 clusters across 13,130 astrocytes in the aggregated dataset (Fig. 1a). Parsing out the full factorial design of the study to examine the distribution of astrocyte cluster subtypes as a function genotype by treatment revealed striking segregation of astrocyte subtypes (Fig. 1b). LPS-repressed subtypes were found to be As_1, As_2, As_3, As_4, As_ 5, As_7, and As_10, with proportional distribution between ApoE3 and ApoE4 genotypes (Fig. 1b/c). Interestingly, LPS-enriched subtypes were found to be As_0, As_6, As_8, As_9, and As_11, which proportionally demonstrate some genotype-specific bias, as As_9 was primarily found in ApoE3, while As_8 and As_11 were primarily observed in ApoE4. Examining the top subtype biomarker genes from only the LPS-enriched subsets via k-means clustering revealed class-level groupings corresponding to each astrocyte subtype. *As_0* had expression profiles associated with extracellular matrix and blood brain barrier modifiers (*Fbln5, Agt, Sparc*) ^18,19^. As_6 was a high-expressor of *Scrg1, Mfge8, S1pr1*, and *Gnao1* which have been linked with clasmatodendrosis/vacuolization, neuronal phagocytosis, neuron-astrocyte cross-talk, and calcium signaling during neuroinflammation, respectively ^20–24^. As_9, over-represented in the E3 LPS brain, exhibited hallmark genes associated with scar formation and wound healing (*Cspg5, Htra1*), ^25,26^, while E4 LPS-enriched As_8/11 subtypes clustered similarly together and expressed genes associated with pro-inflammatory response (*Lcn2, Cxcl10*), as well as oxidative stress and metal cation buffering (Fig. 1d; *Hsbp1, S100a6, Mt1, Mt2*) ^27–29^. Using Monocle2 as an alternate analysis pipeline, we demonstrate via pseudotime trajectory that several of the hallmark genes from As_8/11 are highly expressed in the E4_LPS condition (Fig 1e; purple).

**Figure 1.**
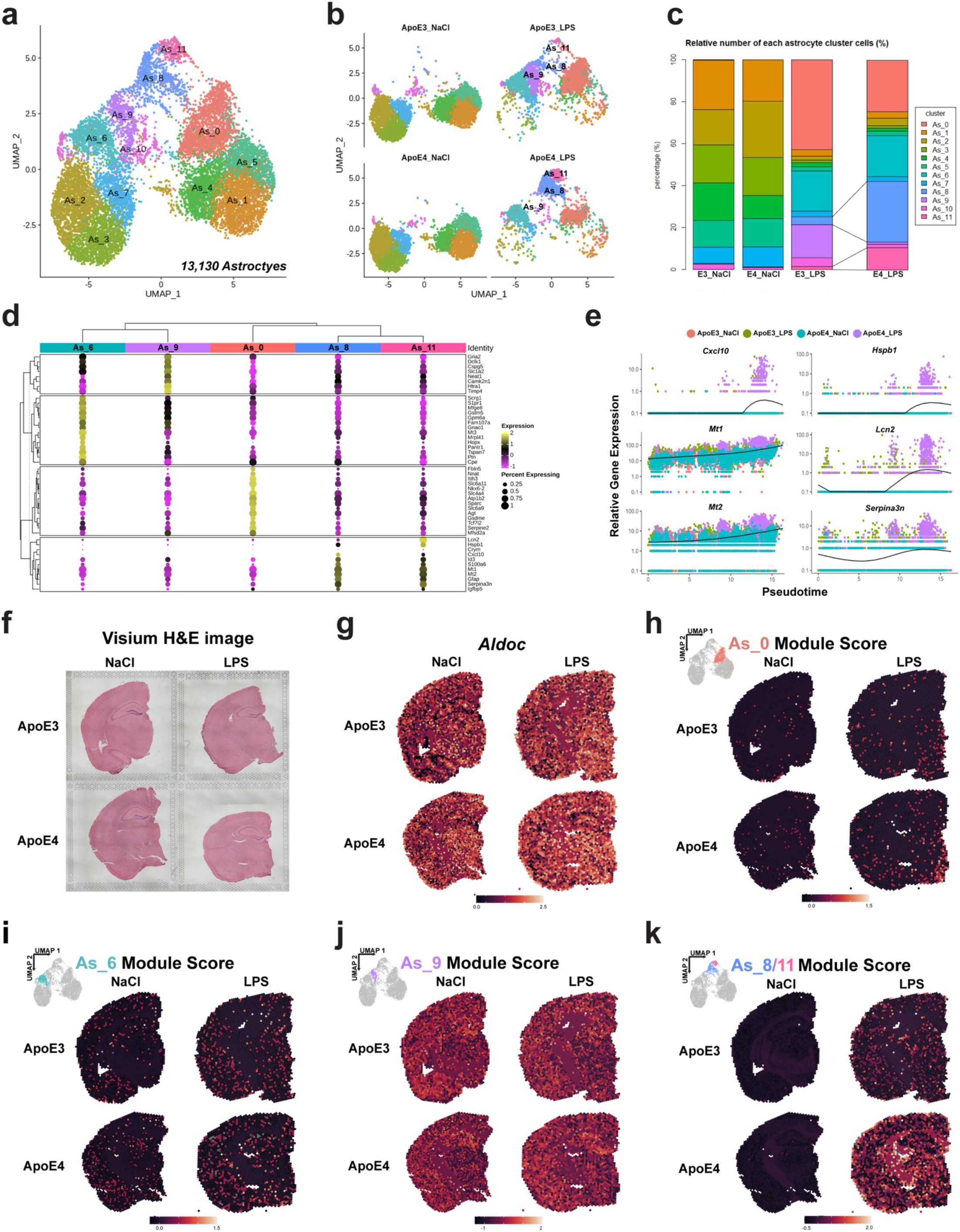
ApoE4 drives transcriptional heterogeneity in astrocytes. **a** 13,130 astrocytes across four pooled samples are visualized via UMAP in the dimensionally reduced dataset, representing 12 subtypes. **b** Splitting of the aggregated dataset by genotype and treatment reveals prominent shift in subtype proportions across the factorial design. Visually, there was a noticeable shift away from seven As_subtypes (1, 2, 3, 4, 5, 7, 10) found predominantly in the NaCl-treated mice and accumulation of LPS-associated subtypes (0, 6, 8, 9, 11). Notably, there were clear shifts in subtype proportions with an E3-bias for accumulating As_9, while E4 had predominantly more As_8 and As_11 subtypes. **c** Calculation of the percentage of each As_subtype by genotype and treatment. **d** k-means clustered (k = 4) dot plot using the top five differentially expressed biomarkers for each of the LPS-enriched astrocyte subtypes. **e** Cell trajectory inference analysis calculations of astrocyte pseudotime score plotted against selected gene expression values of representative As8/11 subsets demonstrating E4-associated bias (teal and purple). **f** H&E brightfield image of the Visium spatial profiling tissue sections. **g** Expression score of *Aldoc*, a gene enriched in astrocytes, shows robust tissue-wide expression across all sections. **h-k** Module score expression visualization utilizing the differentially expressed biomarker genesets for LPS-enriched subtypes As_0, As_6, As_9, and As_8/11, respectively.

Recent reports have demonstrated regionally-associated astrocyte transcriptional profiles ^17,30,31^. From a separate cohort of mice we generated spatial transcriptional datasets to determine whether carriage of either ApoE allele altered the regionally associated transcriptional profiles of our LPS-enriched As_subtypes. We calculated the “module score” utilizing the k-means clustered biomarkers from each LPS-enriched astrocyte subtype against each spot within the aggregated 4 sample spatial dataset. Plotting these module scores for each did not demonstrate a remarkable bias in a particular anatomical region for any cluster (Figs. 1g-k). However, we were able to resolve genotype biases following LPS, similar to our scRNAseq findings, with As_8/11 having increased expression at the spot-level in the E4-LPS condition, relative to E3-LPS. These signatures were notably absent in the NaCl administered groups, irrespective of genotype.

### ApoE4-associated transcriptional alterations in astrocyte states implicate maladaptive immunometabolic phenotypes

We assessed the gene-gene co-expression network of our scRNAseq dataset using the MEGENA pipeline ^32^. Nine of the twelve astrocyte subtypes displayed significant enrichment overlap with the gene-gene network clustering (Fig. 2a). In particular, parent MEGENA-defined network clusters 2 and 3 (c1_2, c1_3) displayed significant enrichment with As_8/11 astrocyte subtypes. Expansion of the gene networks of these subsets revealed several hub genes, including Gfap, *Mt1, Mt2, Fth1, Serpina3n, Lcn2*. Examining the hub genes from cluster 2 (i.e. C1_2) across our spatial dataset revealed enrichment in the ApoE4-LPS tissue, with some regionality apparent in the borders, white matter tracks, and internal capsule (Fig. 2c). Using focused analysis parameters of the LPS-enriched clusters to delineate pathway enrichment of As_8/11 from the other subsets, we show that this E4-enriched subset acquires phenotypes associated with MHCI signaling, interferon response, high translation and ribosomal activity, as well as signaling via NFkB (Fig. 2d). Integrating the NFkB pathway genesets across our spatial landscape revealed that the E4-LPS tissue had similarly enriched expression profiles compared to E3-LPS (Fig. 2e).

**Figure 2.**
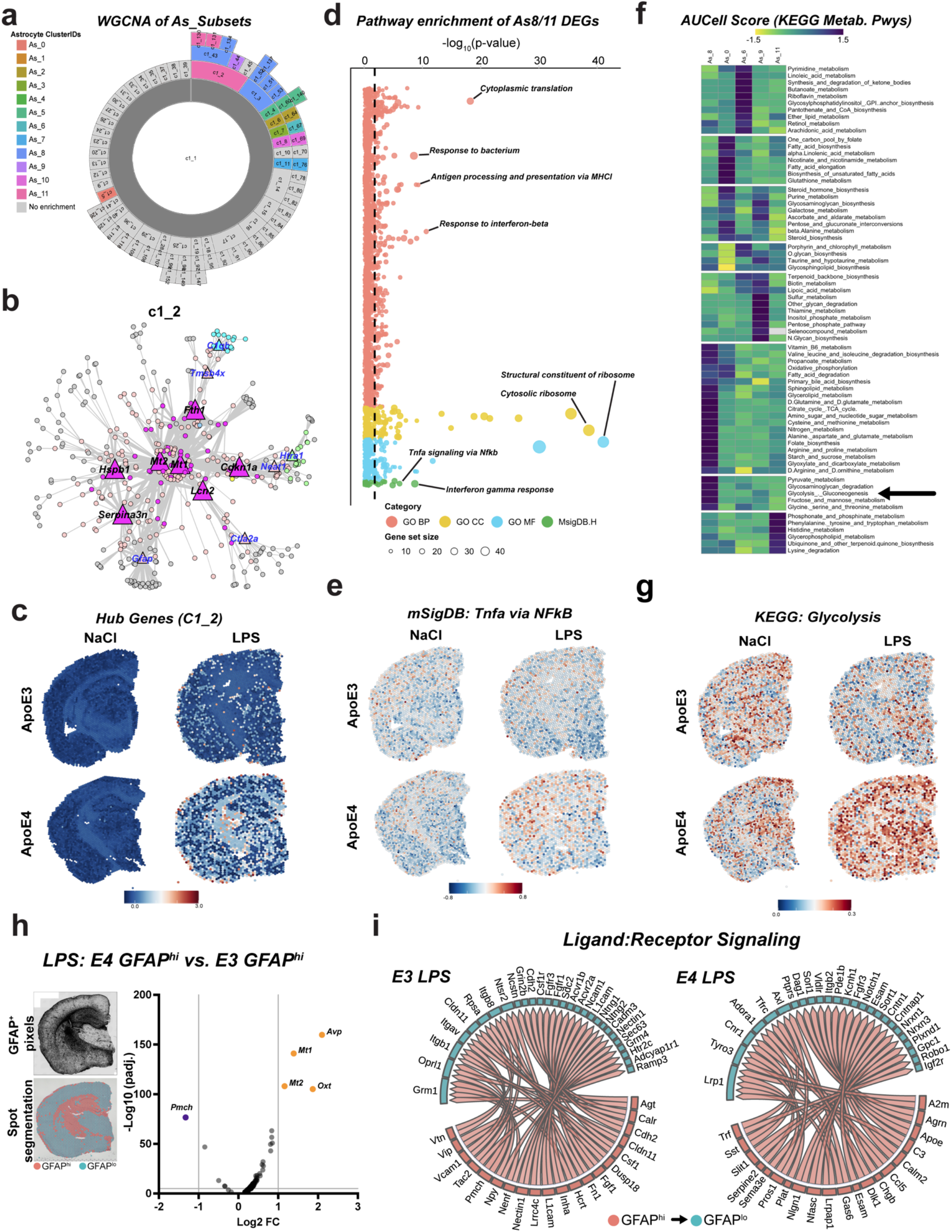
ApoE4 is associated with differential immunometabolic signatures of astrocytes. **a** Sunburst plot of the MEGENA co-expression network hierarchy of the scRNAseq pseudo-bulk expression data. This hierarchical network visualization of concentric rings represents root parent modules that are increasingly subdivided into child modules, moving outward. Modules are colored according to significant overlap (FDR <0.05, Fishers exact test) with As_subtypes. **b** MEGENA module (c1_2) displayed significant overlap with the cell-type enrichment of E4-associated LPS subtypes As_8/11, hub genes (triangles) and their associated networked genes (circles) are indicated wherein there was a significant difference in expression between E4 and E3 astrocytes (FDR <0.05). **c** Integrating these hub genes as an aggregate module score across our spatial datasets demonstrates striking enrichment across multiple anatomical areas in the E4-LPS condition. **d** Pathway enrichment analyses of E4-enriched As_8/11 subtypes demonstrate significant correlations with multiple ontologies related to immune signaling and cellular activity, with representative pathways highlighted for each ontology. **e** Geneset defining the representative As_8/11 enriched ontologies integrated back into spatial datasets for ‘Tnfa-signaling via Nfkb’, showing higher expression across the E4-LPS tissue section, comparatively. **f** KEGG metabolic pathway enrichment analysis via AUCell examining the LPS-associated As_subtypes shows differential alignment across metabolic pathways, with subtypes As_8/11 having marked divergent profiles compared to the E3-LPS enriched As_9 subtype. Glycolysis/Gluconeogenesis has been highlighted (arrow). **g** Genes for KEGG:Glycolysis_gluconeogenesis were analyzed across spatial dataset, which demonstrate an enrichment in the E4-LPS tissue. **h** Log2 fold change from DESeq2 pairwise comparison of E4:LPS vs. E3:LPS through spot segmentation of GFAP^+^ pixels (inset; black and white) with spatial transcripts only from segmented spots (inset; red ‘GFAP^hi^’ spots). **i** NicheNet ligand:receptor interaction Circos plots visualizing putative signaling from GFAP^hi^ segments to GFAP^lo^ in the LPS-treated tissue split by genotype (E3 left, E4 right).

Given consistent association with E4-driven alterations in cellular metabolism ^33,34^, we assessed the alignment of KEGG metabolic pathways across the LPS-enriched astrocyte subsets using AUCell ^35^ to determine whether these neuroinflammatory-responsive subsets acquire preferential enrichment of metabolic transcriptional pathways. Our findings indicate that overall, there was distinct enrichment between the multiple LPS-enriched subsets (Fig. 2f). The E3-LPS predominating subset As_9 showed high expression of pathways related to glycan biosynthesis and metabolism as well as pentose phosphate pathway, while the E4-LPS enriched subsets of As_8/11 had overlapping scores with Oxidative phosphorylation, Glycolysis/Gluconeogenesis, Fatty acid degradation, and Glycerophospholipid pathways. Strikingly, when we integrated the geneset for Glycolysis/Gluconeogenesis into our spatial dataset, we observed a marked increase in this expression module in the E4-LPS tissue (Fig. 2g).

Lastly, we extended our spatial dataset by incorporating protein expression of GFAP histology and correlating its immunofluorescent signal intensities with spot-level gene expression using DESeq2 (E4-LPS vs. E3-LPS). Our findings indicate spots with high levels of GFAP protein expression were also correlated with ‘hub/hallmark’ genes *Mt1* and *Mt2* that were identified in our single cell dataset (Fig. 2h), corroborating that the areas with high GFAP-reactive astrocytes in the E4-LPS condition are linked with these inflammatory genes. Furthermore, when we assessed the putative connectivity via NicheNet across this spatial domain between areas defined with high versus low GFAP intensities (Fig 2h, (inset) red vs. blue spots) we observed differential interactions based upon genotype. Principally, the ligand:receptor pairs in the E3_LPS tissue (Fig. 2i; left) had gene pairs associated with promotion of cerebral blood flow and synaptic function (*Agt*), maintenance of microglia (*Csf1*), as well as mitigation of oxidative stress (*Fgf1*). Comparatively, these areas of high GFAP reactivity in the E4_LPS tissue (Fig. 2i; right) had gene pairs associated with higher lipoprotein signaling/metabolism (*Apoe, Lrp1, Vldr*), TAM receptor signaling/synapse elimination (*Gas6, Axl, Tyro3*), inflammatory response (*Ccl5*), and complement (*C3*), collectively implicating increased astrocyte:microglial cross-talk. Taken together, integration of single-cell and spatial transcriptomics highlights an E4-driven disruption in multiple domains linked astrocyte:microglia inflammatory signaling, synaptic phagocytosis, and cellular metabolism, compared to E3 tissues, which tend to preserve mechanisms associated with neuroprotective/homeostatic response.

### ApoE4 drives altered immunometabolic responses in isogenic iPSC astrocytes

To confirm these findings with functional metabolic indices, we next examined a reduced model system utilizing isogenic E3/E3 and E4/E4 iPSC-derived astrocytes from TCW et al. ^34^. Prior work examining potential inflammatory mechanisms underlying astrocyte response to systemic LPS administration identified TNF, IL1α, and C1q as microglia-derived soluble constituents influencing astrocyte reactivity ^13^. Utilizing this cocktail of pro-inflammatory mediators, we examined cellular respiration and glycolytic response using the Seahorse system. Similar to previous studies in both immortalized and primary murine astrocytes ^33,36,37^, E4-expressing iPSC-derived astrocytes had lower oxygen consumption rates and increased glycolysis compared to E3 cells (Fig 3a; light colors). Interestingly, following the pro-inflammatory challenge, E4-expressing astrocytes further suppressed their maximal respiratory capacity in addition to showing additional increases in both basal and compensatory glycolysis (Fig. 3a; dark red). Plotting the cumulative energetic mapping of both assays demonstrates that while the E3-expressing astrocytes primarily respond to the pro-inflammatory cocktail by increasing mitochondrial-associated respiration, E4 astrocytes conversely respond with a robust increase in glycolysis (Fig. 3b).

**Figure 3.**
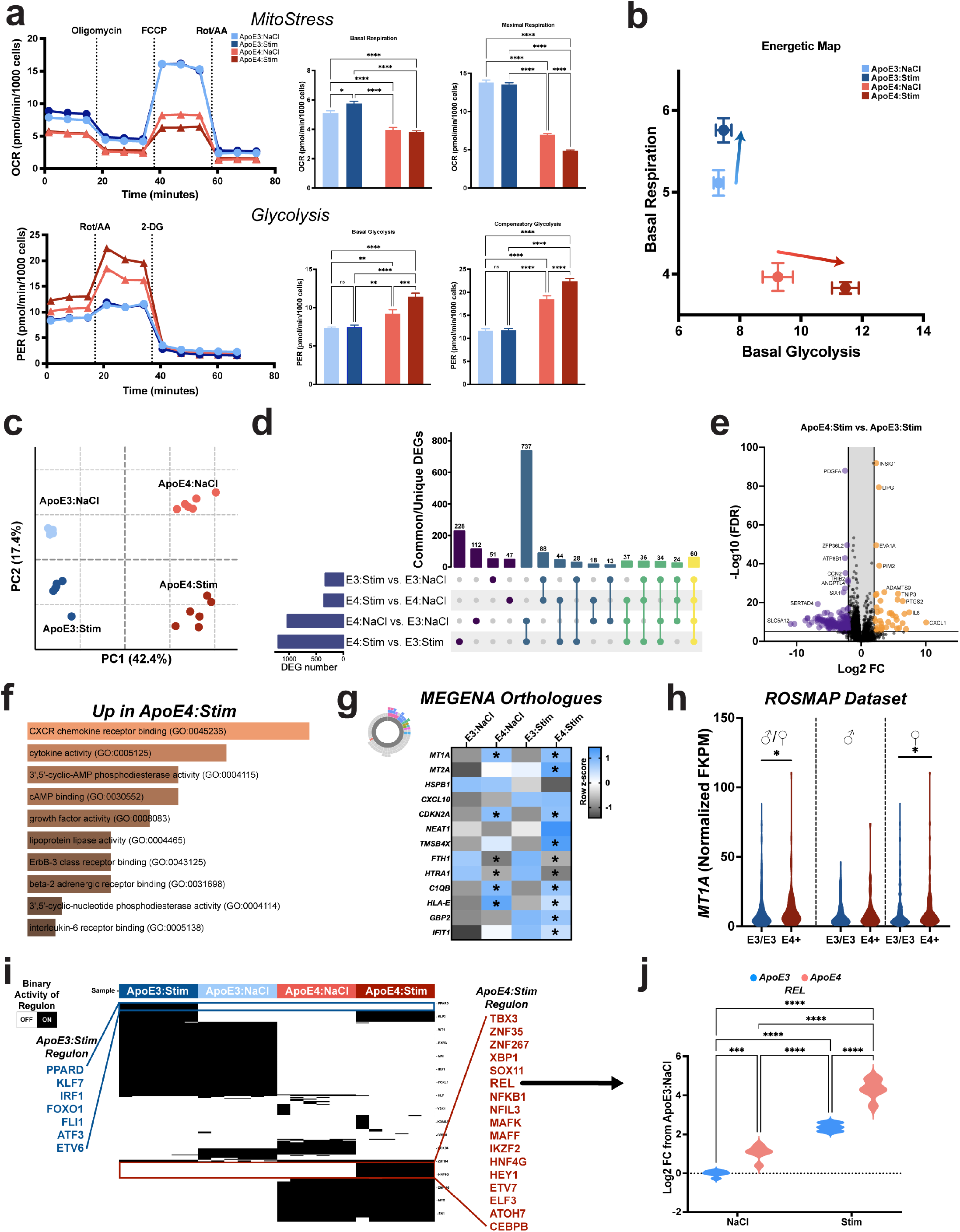
ApoE4 exacerbates pro-inflammatory induced glycolytic shift in human iPSC astrocytes. **a** Mitochondrial respiration and cellular glycolytic capacity were assessed via Seahorse assays for iPSC-derived human astrocytes isogenically encoding either E3/E3 or E4/E4 alleles with or without pro-inflammatory stimulation. Consistently, harboring E4 alleles drove a main effect alterations in basal respiration (2way ANOVA, F(1, 87) = 117.5, p<0.0001) and glycolytic capacity (2way ANOVA; F(1, 66) = 69.10, p<0.0001, compared to E3. Challenging E4 iPSC astrocytes with pro-inflammatory stimulus significantly altered patterns in basal glycolysis (Tukey’s; p = 0.0003), maximal respiration (Tukey’s; p<0.0001), and compensatory glycolysis (Tukey’s; p<0.0001), compared to E4:NaCl (n=20-22 technical replicates per group). **b** Plot demonstrating the aggregate energetic mapping of cell states across basal respiration and glycolysis, with visual shift in E4 astrocytes to move further away from respiration along a primarily glycolytic trajectory. **c** Bulk RNAsequencing from the above four conditions (n=6 per group) plotting the top 5000 most variable features within PCA (69.8% variance explained) showing clear demarcations across genotype (PC1; 42.4%) and treatment (PC2; 17.4%). **d** Upset plot comparing four conditions’ DESeq2 FDR-corrected (>2logFC) pairwise comparisons. Comparatively, there were 228 unique DEGs found within the E4:Stim vs. E3:Stim pairwise contrast. **e** Volcano plot highlighting several up- and downregulated DEGs from E4:Stim vs. E3:Stim contrast. **f** EnrichR pathway ontology analysis of up-regulated DEGs from E4:Stim vs. E3:Stim. **g** As_8/11 MEGENA-defined hub gene human orthologues analyzed using 2way ANOVA with Šídák’s posthoc comparisons examining E4-bias in saline or stimulated responses (*p<0.05 for each corrected pairwise contrast). **h** *MT1A* orthologue quantified from ROSMAP dataset demonstrates E4-biased (E3/E4 and E4/E4) increased expression (Mann-Whitney, p = 0.0375), when disaggregated by sex only female E4+ carriers had a significant increase in expression (Mann-Whitney, p = 0.0189). **i** Transcription factor (TF) inference analysis via PyScenic data binarization predicts 17 unique TF regulons in the E4:Stim condition relative to all. **j** Predicted TF of the *REL* NFkB canonical signaling expressed as Log2FC from E3:NaCl, with both a main effect found due to harboring E4 (2way ANOVA; F(1,20) = 119, p<0.0001) and pairwise comparison relative to E4:Stim (Tukey’s; p<0.0001).

Extending this paradigm, we quantified the transcriptional repertoire of the iPSC-derived astrocytes using bulk sequencing. PCA analysis of the top variable features shows a separation in the orthogonal dataset across PCs visually linking to genotype (PC1, 42.4% variance explained) and treatment (PC2, 17.4% variance explained). This suggests that the transcriptional profile associated with ApoE4 is sufficiently distinct compared to ApoE3 and importantly, their respective responses to pro-inflammatory challenge are divergent (Fig. 3c). We calculated DEGs utilizing DESeq2 for the four pre-planned pairwise contrasts to isolate genotype, treatment, and combined interactions. Plotting the FDR corrected DEGs in an Upset plot to identify both unique and divergent expression profiles revealed there were 228 uniquely enriched DEGs in the ApoE4:Stim vs. ApoE3:Stim pairwise contrast, compared to all other pairwise comparisons (Fig. 3d). Further, the largest proportion of unique DEGs was attributed to differences associated with ApoE4 relative to ApoE3, resulting in 737 linked DEGs between both pairwise comparisons. Volcano plot of the ApoE4:Stim versus ApoE3:Stim pairwise comparsion visualized several of these representative DEGs, which were enriched for genes involved in the pro-inflammatory response (*IL6, CXCL1, TNIP3*) in the E4:Stim condition (Fig. 3e). Pathway analysis of these upregulated genes highlighted pathways such as CXCR1 receptor signaling, cytokine activity, cyclic-AMP activity, lipoprotein lipase response, and IL6 receptor signaling (Fig. 3f), which aligns with previously reported responses in these isogenic cell lines ^34^. This suggests that E4 astrocytes preferentially drive exacerbated pro-inflammatory transcriptional tendencies, compared to E3-expressing astrocytes.

To understand whether the E4-enriched biomarkers we identified in our mouse studies (i.e. Figs 1 & 2) translated over to human-derived iPSC astrocytes, we compared several of the top differentially expressed hub genes identified via MEGENA (e.g. Figs 2a/b). For these selected orthologues, we observed multiple patterns wherein ApoE4 was associated with significant alterations in the expression levels of these genes, up- or downregulated, as compared to ApoE3 iPSC astrocytes. Specifically, *MT1A and MT2A* (i.e. *Mt1/Mt2*) were significantly increased in the E4:Stim vs. E3:Stim (Fig. 3g), with *GBP2, CDKN1A, C1QB, HLA-E* (i.e. *H2-T23*), following a similar expression pattern. *CXCL10* appeared to be a generally upregulated in response to the pro-inflammatory stimulus, but was not significantly different between E3:Stim vs E4:Stim astrocytes. Given that we observed a conserved expression pattern from our APOE4 mice and E4 iPSC astrocytes with *Mt1/MT1A*, we next examined whether this pattern was present in human brain tissue. Utilizing the ROSMAP dataset, we extracted the normalized FKPM values from homozygous E3 carriers and E4 carriers (E3/E4 and E4/E4 were combined). Our findings revealed that in the E4+ carriers, *MT1A* was significantly upregulated compared to E3/E3 carriers (Fig. 3h). When disaggregated for sex, MT1A was significantly increased in the female E4+ carriers with a similar nonsignificant trend in the male E4+ carriers (Fig. 3h).

Lastly, we sought to examine potential underlying transcription factors that would be correlated with the transcriptional expression patterns in our reduced model system. Utilizing the *PyScenic* package – which binarizes activity states into expression regulons - our data reveal several conserved inferences of transcription factor (TF) activity across our four conditions. Interestingly, while the ApoE3:Stim condition revealed differentially predicted TFs associated with inflammatory response (e.g. IRF1 and FLI1), there were also alignment with TFs previously linked with astrocyte-associated anti-inflammatory response via PPAR ^38,39^ as well as KLF7. KLF7 in astrocytes has been linked with axonogenesis ^31^ as well as a mediator promoting Stat3 signaling cascades associated wound repair ^40^. Comparatively, in the E4:Stim condition we observed several TFs associated with regulating inflammatory response (*NFIL3, NFKB1, MAFF, MAFK, CEBPB*) as well as canonical NFkB signaling attributed to *REL* (Fig. 3i). As our single cell dataset had also suggested pathway alignment with NFkB signaling (Fig. 2), we extracted the normalized FKPM expression for *REL* across our four conditions. Expression of *REL* was significantly upregulated following pro-inflammatory challenge in E3 cells, however, this was surpassed in the basal E4 condition and further amplified in the E4:Stim condition (Fig. 3j). Together, these multimodal data suggest that the E4-associated and pro-inflammatory exacerbated responses of iPSC astrocytes may be linked with preferential utilization of upstream mediators associated with canonical NFkB cascades, which are not robustly engaged in the ApoE3 astrocytes during an inflammatory stressor.

### ApoE4 utilization of REL drives maladaptive immunometabolic responses of human iPSC astrocytes

Together, our single cell, spatial, and bulk sequencing datasets demonstrate a consistent link with NFkB signaling that is preferentially upregulated in E4-expressing astrocytes. In particular, our inference analyses predicted that these E4-associated effects may be driven through the canonical NFκB constituent REL. To ascertain whether the basally heightened and pro-inflammatory stimulus exacerbating expression pattern of REL in E4-expressing astrocytes is of functional consequence we utilized IT-603, a small molecule inhibitor of REL nuclear translocation ^41^. We first assessed whether targeting REL activity was sufficient to blunt the observed E4-specific glycolytic shift (Fig. 4a/b). Interestingly, we observed that while suppression of REL activity significantly mitigated both basal- and spare glycolytic function in the E4-expressing astrocytes, E3 expressing astrocytes were minimally altered (Fig. 4a). In the E4 condition, IT-603 was sufficient to suppress the higher basal and compensatory responses both in control and pro-inflammatory environments (Fig. 4b). As the E4 astrocytes exhibited significant alterations in response to REL activity suppression, we examined their underlying cellular profiles using targeted metabolomics. In general, pro-inflammatory stimulation drove heightened production of multiple metabolites, compared to the unstimulated control condition (Fig. 4c). Notably lactate, alanine, citrate, serine, and glycine (Fig. 4c). Furthermore, extracellular lactate concentrations paralleled cellular levels in their responsiveness to REL suppression without any alterations in the available pool of glucose (Fig. 4d). From the same samples as our cellular metabolite and extracellular lactate measurements, we quantified 24 cytokine/chemokine analytes using multiplex high sensitivity ELISA. As expected, the pro-inflammatory stimulus induced the production of the majority of these analytes. However, in the context of REL inhibition, we observed significant reductions across multiple proteins, including CCL2, CXCL10, IL6, and IL1β (Fig. 4e). Collectively, when we assessed these changes in a multivariate space using PCA (PC1 and PC2; 71.6% of total variance), IT-603 alone did not drive noticeable exudate shifts in phenotype. However, when combined with the pro-inflammatory exposure there was a clear shift along both PC orthogonals (Fig. 4f). Further, when we use multifactorial clustering to examine each sample’s lactate release against its inflammatory measures, we observed a strong positive correlation across secreted IL-8(HA), CCL4, CCL2, CCL3, IL1β, and IL6 (Fig. 4g). Overall, the aggregate findings demonstrate strong functional and correlative relationships implicating ApoE4 utilization of canonical NFkB signaling via REL as a driver of maladaptive immunometabolic capacity.

**Figure 4.**
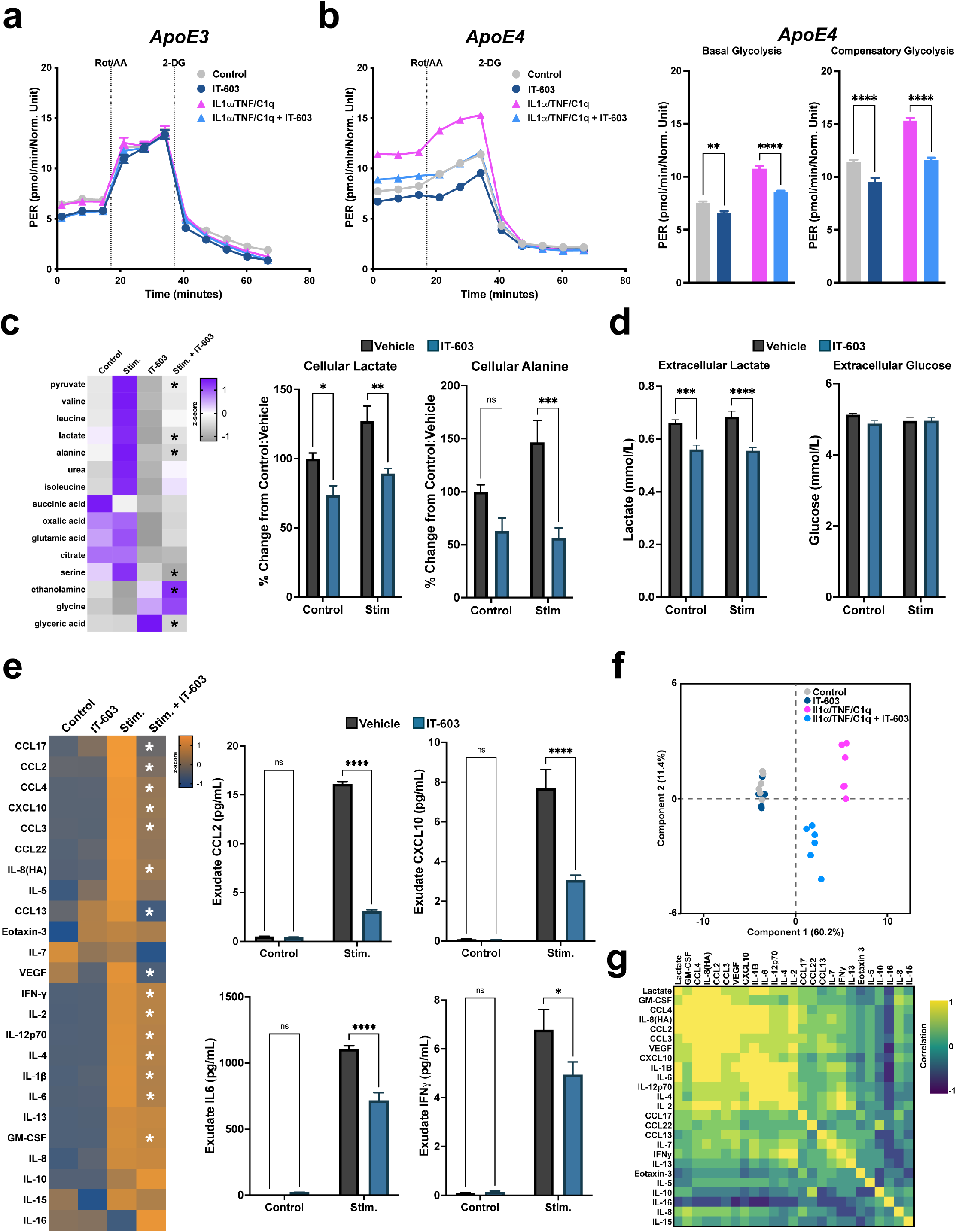
Pharmacological targeting of REL abrogates ApoE4-associated immunometabolic responses in human iPSC astrocytes. **a/b** Inhibition of REL via IT-603 (20μM) had minimal effect on glycolytic response of isogenic E3 astrocytes, but significantly blunted both basal and pro-inflammatory induced exacerbated glycolytic responses of E4 iPSC astrocytes (Šídák’s multiple comparisons posthoc ***p<0.01; n=20-22 technical replicates per group). **c** Cellular steady-state metabolomics revealed IT-603 significantly altered the pro-inflammatory induced alterations in isogenic E4 iPSC astrocytes relative to E4:NaCl baseline (Šídák’s multiple comparisons posthoc *p<0.05; n=5-6 technical replicates per group), representative factorial plots for cellular lactate and alanine production are shown. **d** Quantification of extracellular lactate revealed similar trends as cellular metabolites, with IT-603 driving a significant suppression from vehicle-treated cells (Šídák’s multiple comparisons posthoc *p<0.05; n=5-6 technical replicates per group), but there were no differences in extracellular glucose availability. **e** Multiplex ELISA analysis of exudate cytokine and chemokines revealed consistent upregulation in the majority of soluble analytes, to which IT-603 was sufficient to significantly suppress many of these responses (n= 6/group; Šídák’s multiple comparisons posthoc ***p<0.05). Representative factorial plots for CCL2, CXCL10, IL6, and IFN are shown. **f** PCA plot (71.6% variance explained) of ELISA results shows a clear shift of IT-603 treated E4 iPSC astrocytes away from vehicle treated pro-inflammatory responses, with IT-603-alone causing minimal alteration in the exudate production. **g** Correlation matrix on extracellular lactate vs. ELISA analytes in the E4:Stim vehicle samples, demonstrating clustered enrichment correlation with lactate.

## Discussion

While much of the work investigating CNS immunometabolism in health and disease have predominately focused on microglia, a handful of studies have examined astrocytes. From a metabolic perspective, these studies suggest astrocyte reactivity increases glycolysis while decreasing mitochondrial oxygen consumption, in a similar manner to processes typically associated with cells from innate immune lineages^42–44^.

In the context of ApoE4 astrocyte metabolism, we recently documented that steady-state central carbon metabolism is altered in ApoE4 expressing immortalized astrocytes^33,37^. However, understanding of the molecular mediators underlying ApoE4-driven alterations in immunometabolic response of ApoE4 astrocytes to inflammatory challenge remains obscure. Our results demonstrate a key role for ApoE4 in perpetuating dysfunctional responses to neuroinflammation via (1) disproportionate accumulation of reactive heterogeneity, (2) enhanced pro-inflammatory phenotypes, and (3) altered glycolytic transcriptional and cellular responses, which (4) appear to be mechanistically linked through exacerbated reliance on the canonical NFkB transcription factor REL. These findings further extend and support the notion that ApoE4 regulates astrocyte functionality via pathways associated with maladaptive immunometabolism.

Prior work has consistently demonstrated that ApoE4 astrocytes have altered responses to a variety of neuroinflammatory stimuli, in both acute insults and chronic degenerative cascades ^15,16,45^. Our findings implicate, in part, the maladaptive nature conferred by ApoE4 in astrocytes *in vivo* resides with an imbalance in acquiring inflammatory-driven reactive heterogeneity compared to ApoE3 astrocytes. ApoE4 astrocytes preferentially acquired As_8/11 subsets that were high expressors for genes associated with inflammation, metal cation imbalance and oxidative stress. Specifically, we observed transcriptionally distinct clusters hallmarked by expression of metallothionein gene isoforms *Mt1* and *Mt2*. These have been linked with oxidative stress and metal ion imbalance, to which Mt1 can increase glycolytic output ^46^, while Mt2 can drive cells away from cellular respiration and oxidative phosphorylation ^47^. These genes and their human orthologues were also upregulated in astrocytes sequenced from AD brain tissue ^48^ and have been linked with redox-initiated DNA damage in astrocytes ^49^. Interestingly, metallothioneins were identified to interact with and activate NFkB ^50,51^, suggesting linkage with the increased pro-inflammatory immunometabolic bias we observed. Furthermore, *Mt1* has been demonstrated to negatively regulate the utilization of Stat3 in response to inflammation ^52^. Importantly, Stat3 has been linked with critical resolution and neuroprotective responses of astrocytes to neuroinflammation ^25,53^. Conversely, ApoE3 astrocytes acquired a distinct enrichment of the As_9 subtype, which was minimally observed in ApoE4 animals. This ApoE3-biased response to systemic LPS exposure exhibited hallmark genes with neuroprotective reactive astrocyte modalities associated wound repair, inhibition of matrix metalloproteinases, and calcium balancing. Transcriptionally, our findings point toward an inability of ApoE4-expressing astrocytes to acquire neuroprotective/repair phenotypes that are observed in the E3 condition, and this inflexibility may be rooted in alterations in sensors and pathways associated with metal ion imbalance.

Examining functional correlates associated with acute inflammatory response of ApoE4 expressing astrocytes revealed marked divergence from their isogenic ApoE3 counterparts. Our findings demonstrate that pro-inflammatory challenge via IL1a, C1q, and TNF was sufficient to significantly push E4-expressing astrocytes toward an exacerbated glycolytic response. Interestingly, these responses were in contrast to ApoE3 iPSC astrocytes, which demonstrated minimal glycolytic shift and maintenance of mitochondrial respiration. These functional metabolic measures paralleled/mimicked our transcriptional assessments *in vivo*, which demonstrated an apparent lack of metabolic flexibility in E4 astrocytes in response to neuroinflammation. When we examined the bulk transcriptional response in isogenic human iPSC astrocytes, we observed unique differential enrichment of multiple pro-inflammatory cascades, compared to ApoE3 astrocytes. Importantly, we demonstrate that the human orthologues of gene biomarkers defining As_8/11 were differentially regulated due to *APOE* and/or *APOE* by stimulus interaction, suggesting that our ApoE4-LPS features do not exist solely within a single cell environment. Moreover, we consistently observed *Mt1/MT1A* as an interaction (genotype by neuroinflammation) differentiator between ApoE4 and ApoE3, a finding reflected in human AD brain tissue where E4 carriers showed increased expression of *MT1A* (E4/E4 and E3/E4). Importantly, our findings in terms of increased pro-inflammatory bias for IL6 and CXCL1 expression were in line with the recent report first characterizing these isogenic cell lines ^34^. Here, we extend our current understanding on the underlying mediators of these pro-inflammatory biases is through the use of transcription factor inference analyses by defining multiple unique motifs that are preferentially enriched in ApoE4 astrocytes exposed to acute neuroinflammatory milieu. Recent work using the same isogenic human ApoE3 iPSC astrocytes had demonstrated an inflammatory-induced utilization of canonical NFkB ^54^. Interestingly, our current findings extend these observations by demonstrating that ApoE4 astrocytes preferentially utilize the canonical NFkB Rel transcription factor at baseline, and that there is a synergistic increase in its expression relative to ApoE3 astrocytes in response to inflammatory challenge.

Finally, we sought to understand whether the E4-driven bias across several of our endpoints were functionally linked with Rel. Interestingly, when repeating seahorse glycolytic assays across ApoE3 and ApoE4 iPSC astrocytes, but now in the context of Rel inhibition via IT-603, we did not observe a significant shift in E3-expressing cells. In contrast, we found that inhibition of Rel restored both basal and compensatory glycolytic response *in vitro*. Furthermore, we observed similar effects due to this inhibition upon multiple metabolites, wherein there was a significant suppression in intracellular glycolysis intermediates, including pyruvate, lactate, alanine, serine, and glyceric acid. Further supporting the immunometabolic linkage of Rel with ApoE4, we find that targeting this pathway was sufficient to blunt the release of multiple cytokine/chemokines.

In summary, we demonstrate that ApoE4 confers maladaptive astrocytic responses to acute neuroinflammatory stimuli mediated through disproportionate cellular heterogenic phenotypes and is driven, in part, through over-utilization of Rel. These studies suggest that Rel may be a novel allele-specific target for ApoE4 carriers for restoration of astrocyte-specific cellular responses in the CNS.

## Limitations of the study

There are several limitations of this current study to acknowledge. Notably, we used LPS to model neuroinflammation, which is a poor proxy for virtually all chronic neurodegenerative diseases. Though we have been deliberate about not over-interpreting these findings in terms of implications within chronic diseases of the CNS, we have demonstrated that several of our hallmarks associated with ApoE4 maladaptive bias recapitulate others’ work in both rodent models of- and human tissue from AD. Furthermore, while we were able to mechanistically assess the efficacy of Rel inhibition *in vitro*, we did not demonstrate whether these attributes confer sparing responses *in vivo*. To this end, guided by our findings, future work is needed to ascertain if targeting Rel, either pharmacologically or using genetic models, in the context of ApoE alleles will also provide sparing across modalities associated with immunometabolic dysfunction in neurodegenerative disease.

## Methods

### Animals

All experiments were conducted in accordance with the National Institutes of Health Guide for the Care and Use of Laboratory Animals and were approved by the Institutional Animal Care and Use Committee of the University of Kentucky. Adult (approximately 8-month-old) male APOE3-TR and APOE4-TR mice were used for all experiments. All mice were group housed 4–5 per cage in individually ventilated cages, in environmentally controlled conditions with standard light cycle (14:10 h light to dark cycle at 21°C) and provided food and water ad libitum.

### Lipopolysaccharides injection

Lipopolysaccharides (LPS, *E. coli*, O55:B5, 3×10^6^ EU/mg, Sigma #L2880) was dissolved in 0.9% sterile saline (NaCl: Henry Schein# 63323-186-01). LPS (5mg/kg) or NaCl was administered *in vivo* by intraperitoneal injection. Immediately following injections, mice were returned to their home cages, which were placed halfway upon heating pads set to approximately 37°C. 24 hours following injection, mice were euthanized for tissue- and endpoint-specific assays as described below.

### Brain tissue harvesting for single cell and spatial transcriptomics

At the prescribed interval, mice (n=2 per genotype/treatment) were anesthetized with 5.0% isoflurane before exsanguination and transcardial perfusion with ice-cold Dulbecco phosphate buffered saline (DPBS; Gibco # 14040133). Following perfusion, brain tissues were removed and bisected upon the midline and simultaneously prepared as detailed below.

### Single cell sequencing tissue prep

The right hemisphere was further dissected to divide out the approximate region encompassing AP -.80 to - 2.50mm from bregma. This piece of freshly dissected tissue from each mouse was immediately transferred into gentleMACS C-tube (Miltenyi #130-093-237) containing Adult Brain Dissociation Kit (ADBK) enzymatic digest reagents (Miltenyi #130-107-677) prepared according to manufacturer’s protocol. Tissues were dissociated using the “37C_ABDK” protocol on the gentleMACS Octo Dissociator instrument (Miltenyi #130-095-937) with heaters attached. After tissue digestion, cell suspensions were processed for debris removal following manufacturer’s suggested ABDK protocol. Following completion of this protocol, cell suspensions were pelleted at 300x*g* for 3 min and gently resuspended in 200μL of DPBS + 0.4% bovine serum albumin (Invitrogen #AM2616). Cells were sequentially filtered two more times using Flowmi cell strainers (70μm pore size, Bel-Art #H13680-0040). Cell counts and viability were assessed using AO/PI cell viability dyes (Logos Biosystems #F23001) in tandem with CellDrop automated cell counter (DeNovix). All samples had >90% viable cells, and were diluted according to 10x Genomics suggested concentrations for capturing approximately 10k cells per library.

### Spatial sequencing tissue

The left hemisphere was embedded in Tissue-Tek O.C.T. compound (#4583) in a 10×10×5mm cryomold (Tissue-Tek #4565) and submerged in dry-ice chilled (approximately −70°C) isopentane (Sigma #M32631). Frozen embedded brain tissues were kept on dry ice until being transferred to −80°C storage.

### Single cell capture and scRNA sequencing

For single cell samples; libraries of each pooled (i.e. genotype and treatment) sample were prepped according to 10xGenomics 3’ v3.1 Chromium NextGem Single Cell user guide with 11 cycles of cDNA amplification, after which approximately 25ng of cDNA was used for the fragmentation reaction, and indexing PCR was accomplished using 13 cycles. Library QC and quantitation was performed on a BioAnalyzer 2100 (Agilent) using High Sensitivity chip (Agilent #5067-4627). Sample libraries were pooled and sequenced across a single lane using NovaSeq 6000 S4 platform using 150bp pair-end reads (Novogene, Sacramento CA) and demultiplexed following sequencing.

### Spatial sequencing tissue processing and library preparation

Left hemibrains were sectioned at 10μm, in coronal orientation on a cryostat (Leica). Sections were mounted on Visium slides (10x Genomics #1000184) and kept sealed in airtight container at −80°C until processed for library preparation following manufacturer’s suggested protocol (Visium spatial gene expression user guide RevE, 10xGenomics). Prior to library preparation, mounted sections were fixed using with methanol and hematoxylin and eosin (H&E) counterstained. Images were obtained on a Nikon NiU microscope with Fi3 color camera. Sections were subsequently permeabilized for 18 minutes processed to obtain cDNA sample libraries, according to manufacturer’s suggested protocols. Sample libraries were prepped using 18 cycles of cDNA amplification, after which approximately 100ng of cDNA was used for the fragmentation reaction, and indexing PCR was accomplished using 12 cycles. Library QC and quantitation was performed on a BioAnalyzer 2100 (Agilent) using High Sensitivity chip (Agilent #5067-4627). Sample libraries were pooled and sequenced across a single lane using NovaSeq 6000 S4 platform using 150bp pair-end reads (Novogene, Sacramento CA) and demultiplexed following sequencing.

### Processing of single cell FASTQ files, dimension reduction and cell clustering

Pre-processing of scRNAseq data was accomplished using Cell Ranger (v6.0.2, 10x Genomics), with Illumina files aligned via STAR (2.5.1b) to the genomic sequence (introns and exons) using the mm10 annotation. Standard pre-processing protocols were followed in Cell Ranger to identify cells above background and the resulting filtered gene matrices were utilized in Seurat (v4.0.3) for single cell analyses ^55,56^. Genes that were detected in 3 or more cells were used for analyses. Single cell QC was performed to exclude cells with less than 200 genes or those that exceeded 2500 genes, and greater than 25% mitochondrial genes. Following QC, the four individual samples (i.e. ApoE3_NaCl, ApoE3_LPS, ApoE4_NaCl, and ApoE4_LPS) were merged into a single Seurat object. Data were normalized, scaled, and variable features found using *SCTransform* with default parameters for the merged object. Subsequently, PCA and UMAP were used for two-dimensional visualization of the dimensionally reduced dataset (with the top 30 PCs used, based upon the total variance explained by each). *FindMarkers* function was used to identify the top10 biomarkers for each cell cluster, cells clusters expressing canonical astrocyte-specific genes (i.e. *Aldoc, Slc2a1, Aqp4*) were identified. These cell clusters were subsequently subsetted and stored as a new object, to which the *SCTransform* and dimension reduction, visualization, and biomarker identification methods described above were re-applied. After subsetting and data normalization, three small groups of distinct clusters with differentially expressed genes (DEGs) corresponding to microglia, ependymal cells, and an unclassifiable group were distinguishable from the main astrocyte groups. Following removal of these unidentified cells, there were a total 13,130 cells classified as astrocytes that were utilized for downstream analyses.

### Processing of spatial sequencing FASTQ files, dimension reduction, spatial clustering, and integration

Pre-processing of spatial transcriptomics data was accomplished using Space Ranger (v1.3.0, 10x Genomics), with Illumina files aligned via STAR (2.5.1b) to the genomic sequence (introns and exons) using the mm10 annotation. Default pre-processing protocols were followed in Space Ranger to identify capture areas and align the H&E tissue images with fiducial alignment grids and the resulting filtered gene matrices were utilized in Seurat (v4.0.3) for spatial transcriptional analyses ^55,56^. The four spatial transcriptomic sets (i.e. ApoE3_NaCl, ApoE3_LPS, ApoE4_NaCl, and ApoE4_LPS) were merged into a single Seurat object. Data were normalized, scaled, variable features found, dimensional reduction were completed as described above. Integration of gene sets corresponding to single cell clusters and pathways were calculated using ‘AddModuleScore’ function with default parameters in Seurat. Visualization of the resultant scores were plotted using ‘SpatialPlot’ function with expression scaling standardized across all for visualized samples.

### Integrative analysis of the GFAP image signal intensity and spatial transcriptomic data

Fresh frozen tissue serial sections that were immediately preceding the tissue section used for spatial transcriptomics were used for immunofluorescently labeling GFAP^+^ astrocytes. Briefly, slides containing the adjacent sections to Visium samples were removed from −80°C storage and allowed to acclimate to room temperature in a covered container for 5 minutes. A buffer dam around each tissue section was created with a PAP pen (Vector #H4000). Next, a mixture of acetone/methanol (50:50) stored at −20°C was applied to the tissue for 3 minutes to fix the tissue. Sections were washed with PBS, 3 times, for 10 minutes each. Subsequently, a blocking solution containing 10% goat serum (Lampire #7332500) and 0.2% TX-100 (Sigma) in PBS was applied to the tissue for 1 hour at room temperature. The blocking solution was removed and incubation with GFAP primary antibody (Invitrogen #13-0300, 1:400), diluted in PBS containing 3% goat serum with 0.2% TX-100 (PBST), was applied and sections were incubated overnight at 4°C in a humidity chamber filled with PBS. Sections were washed 3x in PBST, 10 minutes each. Following washes, goat anti-rat secondary (AF568, Invitrogen #A11077, 1:400) diluted in PBST was applied and incubated at room temperature for 2 hours in a humidity chamber filled with PBS. Tissue sections were subsequently washed 3x in PBS for 10 minutes each and allowed to dry overnight at room temperature in a dark box. Slides were coverslipped using mounting media containing DAPI (Prolong Gold, Invitrogen #P36935). Tissue sections were imaged using a Zeiss AxioscanZ.1 with a 20x objective using standardized acquisition parameters, such that all sections were acquired using the same settings. Images of the four tissue sections, in .tiff format, were manually aligned using the DAPI image to the H&E brightfield images from the SpaceRanger output (e.g. ‘tissue_hires_image.png’) acquired for the adjacent Visium sections, in Photoshop (Adobe, v22.4.3). Section alignment was achieved by setting the opacity of the DAPI+ image to 50% and overlaying it above the H&E image as a second layer in Photoshop. Anatomical hallmarks of nuclei staining in the dentate gyrus, CA1, CA3, as well as tissue edges and ventricles were used to align the images and their labeled nuclei as precisely as possible to each other. The aligned imaged containing only GFAP+ RGB values was exported as a .tiff and used for integrating spot-level GFAP signal intensity via Squidpy (v1.0.0) pipeline ^57^. The resulting GFAP intensity scores were added as spot-level metadata. Pairwise comparisons for differentially expressed genes was accomplished using DESeq2 ^58^, while ligand receptor interations between GFAPhi and GFAPlo were generated using NicheNet ^59^.

### Gene co-expression (module) network analyses using MEGENA

Gene co-expression module and hub-gene identification analysis were performed separately for astrocyte populations using the MEGENA (v1.4.1) R software package ^32^. The top 3,000 most variably expressed genes were selected as input. The MEGENA pipeline was then applied using default parameters, using Pearson’s correlations, and minimum module size of 10 genes. Parent modules were produced from which a sub-set of genes form smaller child modules. Modules with > 10 genes were retained for downstream analysis, interpretation, and presentation of results. Co-expression modules were represented graphically using Cytoscape software (v.3.8.2), with hub genes represented with a triangle and nodes with a circle proportional to the node degree.

### Gene set enrichment analyses

The DEG analysis was conducted by the Seurat function FindMarkers via grouping for comparison by clusters, genotype, or treatment (min.pct was set as 0.25 and logFC.threshold was set as 0.25). The DEGs were selected if the adjusted p-value was less than 0.05 and the absolute value of log-fold change was higher than 0.25. Based on the identified DEGs, the enrichment analyses of GO terms (i.e. Molecular Function; MF) were performed via the clusterProfiler R package ^60^. The enrichment analysis results were filtered out if the adjusted p-value was greater than 0.01.

### Gene transcriptional regulatory network analyses using pySCENIC

For regulon identification, gene regulatory network analysis was performed using the pySCENIC software packages (v.0.11.2) ^61^. The arboreto package is used for this step using algorithm of GRNBoost2 (version 0.11.2) to identify potential transcriptional factor (TF)-targets based on their co-expression with RcisTarget (version 1.12.0) for cis-regulatory motif enrichment analysis in the promoter of target genes (mm9-500bp-upstream-10species.mc9nr and mm9-tss-centered-10kb-10species.mc9nr databases provided in the pySCENIC package) and identify the regulon, which consists of a TF and its co-expressed target genes. Correlations between a list of 1,390 human transcription factors (TFs) curated by Lambert et al. ^62^ and the genes in the expression matrix were evaluated, and co-expression modules with a minimum size of 20 genes were defined. Finally, for each regulon, pySCENIC uses the AUCell algorithm to score the regulon activity in each cell. The input for SCENIC was the n (genes) by n (cells) matrix obtained after filtering, and gene expression is reported in count units. Parameters used for running were specified as default options in the original pySCENIC pipeline. The cellular activity pattern of a predicted regulon can be binarized as being in an ‘on’ or ‘off’ state based on the bimodal distribution of a regulon’s AUCell values and visualized as a heatmap for identification of the regulon clusterings.

### Trajectory and pseudotime analysis

To infer the pseudotime of astrocytes progression toward response development, we used functions provided with the Monocle 2 package (version 2.22.0). Pseudotime measures how much progress an individual cell has made through a process such as a cell differentiation or transformation. Seurat data objects were reformatted using the SingleCellExperiment package data format for pseudotime and trajectory analysis sing monocle 2.

### iPSC astrocyte cell culture

Isogenic APOE3 and APOE4 astrocytes ^34^ were initiated at 1×10^6^ cells per mL in complete astrocyte medium (ScienCell #1801) containing 2% FBS (VWR #97068-085) and 1% pen/strep (Gibco #15140-122) and 1% astrocyte growth supplement (ScienCell #1852) plated into a single well of a 6 well dish that was precoated with Matrigel solution (.4% dissolved in complete astrocyte medium; ScienCell #254277) and grown for two days at 37°C, 5% CO2. After this growth phase, cells were passaged into 100mm cell culture dishes (Alkali Scientific #TD0100) and maintained until confluency.

### Harvesting RNA for bulk RNAseq from stimulated APOE3 and APOE4 iPSC astrocytes

12 well plates (VWR #10062-894) were coated with Matrigel solution, APOE3 and APOE4 astrocytes were seeded at approximately 2.18×10^5^ cells/mL/well and incubated for 23hrs as previously described. Following incubation, medium from each well was replaced with either Stimulus or Control media, as above. After 4hrs, cells were gently harvested following a 10 minute incubation with accutase, transferred to a 2mL screwcap tube and centrifuged at 1000xg for 5 minutes at 4°C. Supernatant was aspirated and the resultant cell pellet was snap frozen. Frozen pellets were lysed for RNA extraction using 350uL RLT Plus Buffer (Qiagen # 74004) containing 1% beta-mercaptoethanol (Sigma# 63689) while the pellet thawed on ice for 20 minutes, each sample was thoroughly mixed by pipette and vortexed prior to proceeding to RNA extraction. RNA was extracted following manufacturer’s suggested protocol (Qiagen #74034), resultant RNA was quantified and QC’d using BioAnalyzer 2100 RNA pico chips (Agilent #5067-1513). Average RIN scores for all samples was 9.825, with the lowest RIN score calculated at 9.4, analysis details described below.

### Bulk RNAseq analysis

Bulk RNA sequencing was performed by Novogene (Sacremento, CA) NovaSeq PE150, for all 24 samples generated (n=6 technical replicates per condition), with an average read depth of 27.25M reads per sample. Reads were aligned using STAR 2.7.8a and quantified using the hg38 annotation reference (RefSeq Transcripts 99 - 2021-08-02). Of the 17,366 genes that were quantified to the hg38 annotation reference, only 15,171 genes were utilized for subsequent analyses following filtering to remove genes that exhibited <=10.0 counts per million as the maximum across the 24 sample dataset. This filtered dataset was utilized as described below for individual analyses.

### ROSMAP data analysis

We obtained bulk brain RNA-seq data and associated metadata, including APOE genotypes, from the Religious Orders Study and Memory and Aging Project (ROSMAP) dataset (https://www.synapse.org/#!Synapse:syn3219045). RNA-seq data from the dorsolateral prefrontal cortex (DLPC) were normalized to fragments per kilobase of transcript per million mapped reads (FPKM) ^63^. We compared APOE4 positive carriers (APOE3/E4 and APOE4/E4) to APOE3/E3 homozygotes to assess gene expression as a factor of APOE4 status. Since expression of some genes differed with sex, males and females were analyzed separately. We performed a Mann Whitney non-parametric test for statistical significance because the expression data were not normally distributed. Nominal p-values are reported.

### Seahorse MitoStress and Glycolysis assays

Seahorse plates (Agilent #101085-004) were precoated with Matrigel solution and left to solidify for one hour at room temperature in cell culture hood. APOE3 and APOE4 astrocytes were harvested from 100mm cell culture dishes using 3mL of accutase (Sigma #A6964) for approximately 10 minutes at which point lifting was visibly noticeable. 2mL of complete astrocyte medium was added to each dish. Cell pellets were gently resuspended in 1mL complete medium and counted using hemocytometer. Wells of the Matrigel-coated plates were setup as follows: sterile DPBS (Gibco#114040-133) was added to columns 1 and 12, with the four corner wells (i.e. A1, H1, A12, H12) left empty. The remaining wells were seeded with 40uL at 20,000 cells/well divided equally between the two genotypes. The seeded plates were returned to the incubate for approximately 23hrs at 37°C, 5% CO2. Following this period, a 10x pro-inflammatory cocktail ^13^ was made by combining 5mg/mL IL-1a stock (Sigma #I3901), 10mg/mL TNF stock (Cell Signaling #8902SF), and 1mg/mL C1q stock (MyBioSource #MBS147305) sterile DPBS. The 10x cocktail was subsequently diluted in complete astrocyte medium to generate a 1x pro-inflammatory “Stimulus” of 3ng/mL IL-1a, 30ng/mL TNF, and 400ng/mL C1q. “Control” medium was created by adding the same volume of sterile DPBS into complete astrocyte medium. Medium of each seeded well was gently aspirated and replaced with either “Stimulus” or “Control” medium and incubated for 4hrs at 37°C, 5% CO2. Following incubation, plates were assayed following manufacturer’s suggested protocols for MitoStress (Agilent #103015-100) and Glycolysis Stress (Agilent #103020-100) assays. For experiments examining cRel inhibition, diluted IT-603 (Sigma #5.30654, in DMSO) was added to group-appropriate complete media at a working concentration of 20μM, control groups received the same volume of DMSO alone.

### Harvesting polar metabolites from iPSC isogenic astrocytes for GCMS analysis

Cells were handled in identical manor as described above for bulk RNAseq experiment, with the exception of plating in 6-well dishes at approximately 4.36 × 10^5^ cells per well. Four hours following the media change (stim or control), polar metabolites were extracted for GCMS analysis. Immediately following the last incubatory period of each experiment, plates were removed from the incubator and placed on ice, the media was removed, cells were washed twice with ice cold 0.9% NaCl (VWR BDH Analytical #BDH9286-2.5KG), then 1mL of ice cold 50% MeOH [HPLC-grade (Sigma-Aldrich #A456-4); containing 20μM L-norleucine (Sigma-Aldrich #N6877-1G)] was added to quench cellular metabolic activity followed by 10 min incubation at −80 °C to ensure cell lysis. After removing from the freezer, plates were placed on ice, wells scraped with a cell scraper and the entire contents collected into a tube, vortexed briefly and placed on ice until all samples were collected. Tubes were then placed on a Disruptor Genie Cell Disruptor Homogenizer (Scientific Industries) for 5 min at 3,000 rpm, followed by centrifugation at 24,000 × *g* for 10 min at 4 °C. The aqueous fraction was isolated to a new tube for further processing, the resulting pellet was briefly dried at 10^−3^ mbar CentriVap vacuum concentrator (LabConco) to evaporate any remaining methanol. Following drying, the protein content of the pellet was determined using a BCA assay (ThermoFisher #23225) to normalize metabolite concentrations to total protein amount. The aqueous fraction containing polar metabolites was dried at 10^−3^ mbar followed by derivatization. The dried polar metabolite pellet was derivatized by a two-step methoxyamine protocol first by addition of 50 μL methoxyamine HCl (Sigma-Aldrich #226904-5G) in pyridine (20 mg/mL; Sigma-Aldrich #TS25730) followed by 90 min dry heat incubation at 30 °C. Samples were then centrifuged at 20,000 × g for 10 minutes after which 50μL of each sample was transferred to an amber V-shaped glass chromatography vial (Agilent #5184-3554) containing 80μL N-methyl-trimethylsilyl-trifluoroacetamide (MSTFA; ThermoFisher #TS48915) and gently vortexed followed by 30 min dry heat incubation at 37°C. The samples were allowed to cool to room temperature, and then analyzed via gas chromatography (GC) mass spectrometry (MS).

Samples were analyzed on an Agilent 8890 GC / 5977B MS (Agilent Technologies, Santa Clara, CA, USA). A GC temperature gradient of 130°C was held for 4 min, rising at 6°C/min to 243°C, rising at 60°C/min to 280°C and held for 2 min. Electron ionization energy was set to 70eV. Scan mode for m/z: 50–550 was used for steady-state metabolomics and scan mode for m/z: 50-800 was used for stable-isotope resolved metabolomics. Spectra were translated to relative abundance using the Automated Mass Spectral Deconvolution and Identification System (AMDIS) v2.73 software with retention time and fragmentation pattern matched to FiehnLib library1 ^64^ with a confidence score of > 80.

### Quantification of iPSC astrocyte cellular exudate

Media harvest prior to polar metabolite extraction (above) was centrifuged at 10,000x*g* for 5 minutes at 4°C. Following centrifugation, media for each well was aliquoted and stored at −80°C until analysis. Cytokine and chemokine were quantified using MesoScale Discovery V-Plex Human Cytokine multiplex assays following manufacturer’s instructions (MSD# K15054-D1). Analytes that had ≥50% failure to detect above manufacturer’s lower limit of detection (LLOD) for any one group were omitted entirely from the study, resulting in a total of 24 analytes used for assessment out of the 30 analytes available on the assay. For extracellular lactate and glucose measurements, 20uL from a separate and fresh aliquot per well were thawed on ice and assayed for lactate and glucose concentrations (mmol/L) using a YSI2900 analyzer (YSI incorporated) using the lactate- and glucose-oxidase method per manufacturer’s instructions, as previously described ^65^.

## Acknowledgements

We thank Jim Begley at the University of Kentucky Arts & Sciences Imaging Center for their invaluable assistance with spatial sequencing sample prep. Finally, we thank Dr. Tomoko Sengoku and Michael Alstott at the University of Kentucky Redox Metabolism Shared Resource Facility, supported by NCI Cancer Center Support Grant (P30 CA177558), for their assistance with the Seahorse assays.

## Author Contributions

JMM and LAJ designed the experiments. SL, HCW, LAJ and JMM analyzed the data and wrote the paper. HCW and NAD completed the metabolic analyses of astrocytes, including metabolomics, Seahorse assays. AAG and AEW performed all in vitro experiments. DAH assisted with scRNAseq and spatial seq experiments. DSG and JLS assisted with in vivo experiments, mouse tissue preparation, and ELISA assays. DJZ and SE assisted with ROSMAP analyses. TT and JTCW assisted with iPSC experiments. All authors have read the manuscript and provided edits.

## Data availability

All scRNA-seq and spatial sequencing, and bulk sequencing datasets generated in association with this study are available in the Gene Expression Omnibus under accession number GSE215447. Web-based data browsers are deployed on shiny.servers for both the scRNAseq dataset (https://morgantilab.shinyapps.io/APOEAstroexplorer/) and human iPSC astrocyte bulkseq dataset (https://morgantilab.shinyapps.io/APOExLPS_Bulk).

## Code availability

All codes generated during this study are available upon reasonable request from the corresponding author.

